# Sex differences in fear regulation and reward seeking behaviors in a fear-safety-reward discrimination task

**DOI:** 10.1101/390377

**Authors:** Eliza M. Greiner, Makenzie R. Norris, Iris Müller, Ka H. Ng, Susan Sangha

## Abstract

Reward availability and the potential for danger or safety potently regulate emotion. Despite women being more likely than men to develop emotion dysregulation disorders, there are comparatively few studies investigating fear, safety and reward regulation in females. Here, we show that female Long Evans rats do not suppress conditioned freezing in the presence of a safety cue, nor do they extinguish their freezing response, whereas males do both. Females were also more reward responsive during the reward cue until the first footshock exposure, at which point there were no sex differences in reward seeking to the reward cue. Darting analyses suggest females are able to regulate this behavior in response to the safety cue, suggesting they might be able to discriminate between fear and safety cues but do not demonstrate this with conditioned suppression of freezing behavior. However, levels of darting in this study were too low to make any definitive conclusions. In summary, females showed a significantly different behavioral profile than males in a task that tests the ability to discriminate among fear, safety and reward cues. This paradigm offers a great opportunity to test for mechanisms that are generating these behavioral sex differences in learned safety and reward seeking.

## 1. Introduction

Clinical disorders arising from maladaptive emotion regulation present a large burden on society worldwide. Many of these disorders show comorbidity, for example, addiction with anxiety disorders (Grant et al., 2016). Cues predicting something aversive elicit avoidance and fear behaviors whereas cues predicting reward elicit approach and reward-seeking behaviors. Cues signifying safety have the power to modulate fear and reward-seeking behaviors by informing the organism whether or not the environment is safe (Walasek, Wesierska, & Zieliński, 1995). Thus, safety, fear and reward behaviors, and the circuitries governing these behaviors, are intertwined. The majority of studies on reward and fear processing have been conducted in parallel, investigating the circuitries separately in primarily male subjects. If we hope to understand and treat comorbid disorders resulting from maladaptive emotion regulation, increased efforts in investigating how these circuitries integrate their functions to influence behavior is needed in both male and female subjects.

Our laboratory has designed and validated a behavioral task in which fear, safety and reward cues are learned within the same session allowing us to assess the animal’s ability to discriminate among these cues (Müller, Brinkman, Sowinski, & Sangha, 2018; Ng, Pollock, Urbanczyk, & Sangha, 2018; Sangha, Chadick, & Janak, 2013; Sangha, Greba, Robinson, Ballendine, & Howland, 2014; Sangha, Robinson, Greba, Davies, & Howland, 2014). Rats are exposed to cues associated with safety, fear (fear cue paired with footshock), and reward (reward cue paired with sucrose). Male rats consistently learn to discriminate among safety, fear and reward cues to 1) suppress conditioned freezing in the presence of a safety cue (fear+safety cue), and 2) increase reward seeking when reward is available (reward cue) (Müller et al., 2018; Ng et al., 2018; Sangha et al., 2013; Sangha, Greba, et al., 2014; Sangha, Robinson, et al., 2014). This paradigm also allows us to investigate how safety cues can regulate both fear and reward behaviors. Evidence suggests that reward learning mechanisms overlap at least partially with safety learning (Leknes et al., 2011; Pollak et al., 2008; Rescorla, 1969; Rogan et al., 2005; Sangha et al., 2013; Tanimoto et al., 2004; Walasek et al., 1995). For example, learned safety can act as a behavioral antidepressant in mice (Pollak et al., 2008), and animals will perform certain behaviors in order to turn on a safety signal (Rescorla, 1969; Rogan et al., 2005). Within the amygdala we have shown a subpopulation of neurons responding with the same level of excitation or inhibition during both the reward and safety cues (Sangha et al., 2013). We have also shown a dissociation between reward and safety discrimination; inactivation of the prelimbic or infralimbic cortices of the ventromedial prefrontal cortex have differential effects on reward and safety discrimination, respectively (Sangha, Robinson, et al., 2014). Thus, in male rats, our prior work has already shown a critical involvement of the corticoamygdalar circuit in learning this fear-safety-reward cue discrimination.

Much of the research investigating emotion regulation mechanisms have exclusively used male subjects. In a study using male Vietnam veterans, Post-Traumatic Stress Disorder (PTSD) patients show impairments in suppressing their fear response in the presence of a safety cue (Jovanovic et al., 2009). But, women are more than twice as likely to develop Post-Traumatic Stress Disorder (PTSD) than men, with females having a lifetime prevalence of 8.5% in contrast to 3.4% in males (Mclean et al., 2011). In fear studies that have included female rats, it has been shown that females exhibit lower levels of freezing behavior than male rats after repeated fear cue presentations (Daviu et al., 2014). These findings have been thought to indicate a difficulty in fear conditioning in female rats. A more recent experiment has identified that approximately 40% of female rats tested exhibit an alternate fear behavior in the form of fast paced movements called ‘darting’; this was only seen in approximately 10% of male rats tested (Gruene et al., 2015). There is also evidence of sex differences in the seeking of natural rewards, where it has been reported that female rats consume more sucrose pellets than males and are willing to work harder for them (Tapia, Lee, Weise, Tamasi, & Will, 2019). Dopamine signaling during reward tasks has also been demonstrated to be different between sexes. For example, Conway et al (2019) showed females continue to perform intracranial self-stimulation for brain stimulation reward while under the influence of a kappa-opioid receptor agonist, which suppresses dopamine release, whereas males decrease this behavior. Their data suggest that female rats may have an increased capacity to produce and release dopamine compared to males, under these conditions. Our prior work has shown, in males, that dopamine signaling in the basolateral amygdala contributes to effective discrimination among fear, safety and reward cues (Ng et al., 2018).

Taken together, we hypothesized there would be sex differences in the ability to express clear discrimination among fear, safety and reward cues. The inability of male PTSD patients to learn safety signaling has been labeled a biomarker of the disorder (Jovanovic et al., 2012). Due to sex-related differences in human diagnosis of PTSD, with women diagnosed at rates twice that of men (Glover et al., 2015), any differences female rats have in the learning or retention of safety signals could steer towards further research on the neurological processes underlying these variations.

## 2. Materials and Methods

### 2.1 Subjects

A total of 24 adult male (215-375g) and 28 adult age-matched female (198-230g) Long Evans rats (Blue Spruce; Envigo, Indianapolis), were single-housed and handled for 1 week prior to testing. All procedures were performed during the light cycle and approved by the Purdue Animal Care and Use Committee. Rats had *ad libitum* access to food and water prior to the start of the experiment. After experiment onset, they were maintained on a food restricted diet (20g per day for males; 16g per day for females) until the last day of the experiment.

### 2.2 Apparatus

The rats were trained in operant conditioning chambers consisting of Plexiglas boxes (32cm length x 25cm width x 30cm height) encased in sound-attenuating chambers (Med Associates, ST Albans, VT). 10% liquid sucrose was delivered through a recessed port 2cm above the floor in the center of one wall. Two lights (28V, 100mA) were located 10.5cm from floor on either side of the port. A light (28V, 100mA) 27cm above the floor on the wall opposite the port was on throughout the entire session. Auditory cues were delivered via a speaker (ENV-224BM) located 24cm from the floor on the same wall as the port. Footshocks were delivered through a grid floor via a constant current aversive stimulator (ENV-414S). An overhead video camera and side video camera recorded the sessions for subsequent offline video scoring.

### 2.3 Behavioral Procedures

#### Reward pre-training (5 sessions)

An auditory cue was paired with 10% sucrose solution delivery (100µl) and served as the reward cue (25 trials; ITI, 90-130s).

#### Habituation (1 session)

Rats continued to receive 25 reward cue-sucrose pairings (ITI, 90-130s) in addition to 5 unreinforced presentations each of the future fear and safety cues in order to habituate the rats to their presentation, thereby reducing any baseline freezing to these novel cues.

#### Discriminative conditioning (DC) (4 sessions)

Reward cue-sucrose pairings continued (15 trials). Another auditory cue was paired with a mild 0.5mA, 0.5s footshock and served as the fear cue (4 trials). In separate trials the 20s fear cue was presented at the same time as a 20s safety light cue resulting in no footshock (‘fear+safety’, 15 trials). Trials in which the safety cue was presented alone without any footshock were also included to assess whether freezing developed to the safety cue as well as providing the animal with additional trials that contained a safety cue-no shock contingency (10 trials). Trials were presented pseudorandomly (ITI, 100-140 s). Cues were counterbalanced across subjects with the caveat that the fear+safety compound cue was composed of one auditory cue and one light cue. Eight of the male rats and 12 of the female rats underwent DC training in which the reward cue was a continuous auditory cue (3 kHz, 20s cue; 70dB), the fear cue a pulsing auditory cue (11 kHz, 20s; 70dB), and the safety cue was the presentation of two lights (28V, 100mA located on both sides of the port). The remaining eight male rats and eight female rats underwent training in which the fear and safety cue stimuli were counterbalanced: the light served as the fear cue and the pulsing auditory cue served as the safety cue.

#### Extinction Training (1 session)

One day after the last DC session, both the reward cue and fear cue were presented 20 times each in a pseudorandomized order without sucrose or footshock (ITI, 60-120s).

#### Extinction Test (1 session)

One day after extinction training, rats were presented with the reward (10 trials), fear (10 trials), fear+safety (5 trials) and safety (5 trials) cues in a pseudorandomized order (ITI, 60-120s). None of the cues were presented with sucrose or footshock.

To exclude possible sex differences in pain sensitivity and footshock perception, a separate group of male (n=8) and age-matched female (n=8) rats was presented with a series of unsignalled footshocks of increasing intensities (0.3 mA, 0.35 mA, 0.4 mA, 0.45 mA, 0.5 mA, 0.55 mA, 0.6 mA, 0.7 mA, 0.8 mA, 0.9, 1.0 mA) with an inter-stimulus interval of 2 min. The session was flanked with 5 min intervals in which no stimuli occurred.

### 2.4 Data analyses

Our experimental groups to directly compare males and females on discrimination behavior consisted of 16-20 rats. Cohorts of 4 or 8 female rats were trained alongside cohorts of 4 male rats for a total of 4 replications. Fear behavior was assessed manually offline from videos by measuring freezing, defined as complete immobility with the exception of respiratory movements, which is an innate defensive behavior (Blanchard & Blanchard, 1969; Fendt & Fanselow, 1999). The total time spent freezing during each 20s cue was quantified and expressed as a percentage. Measuring the total time the animal spent inside the reward port and at the entrance of the port with nose positioned at port entrance during each cue assessed reward-seeking behavior and was expressed as a percentage. Darting behavior was detected and quantified offline from videos recorded from overhead cameras via a custom MatLab program, with movements of a velocity of 23.5cm/s or faster qualifying as a single dart (Gruene et al., 2015); these were also confirmed manually. Darting was expressed as the averaged # of darts per cue (sum of darts/ # trials) or trial (sum of darts). Since there were different number of trials per reward, fear, fear+safety and safety cue in each DC session and test for extinction, this was expressed as the sum of darts across trials divided by the number of trials for each cue (sum of darts/ # trials). And, since the extinction training data were expressed trial by trial, data for each individual trial was shown and expressed as the averaged sum of darts for each individual trial (sum of darts). Three individuals performed manual offline behavioral scoring. Pearson’s correlations of behavioral values between scorers were greater than *r* = 0.80. Behavioral data were analyzed with one-way or two-way repeated measures ANOVAs, with sex as the independent factor and condition as the repeated factor, followed by *post hoc* Sidak’s, Tukey’s or Dunnett’s multiple comparisons tests with GraphPad Prism 8. P values were adjusted for multiple comparisons.

For shock sensitivity testing, freezing duration in the 2-min intervals between shock presentations was scored manually, as well as darting and jumping immediately after shock delivery. For the freezing durations, a two-way repeated measures ANOVA was carried out via GraphPad Prism 7, with sex as the independent factor and shock intensity as the repeated factor. Darting and jumping were assessed as dichotomous variables with darting/no darting and jumping/no jumping, respectively. For both, a Cochran test was performed.

## 3. Results

### 3.1 Female rats spent more time reward seeking during reward pre-training

All rats first underwent 5 reward pre-training sessions in which the reward cue was paired with sucrose delivery. The percent time spent at or in the reward port during each reward cue across each reward session was quantified (Figure 1B). Two-way repeated-measures ANOVAs showed main effects of session (F(4,136)=5.395, p=0.0005) and sex (F(1,34)=10.83, p=.0023), but no significant interaction (F(4,136)=0.9031, p=0.4641). *Post hoc* Sidak’s multiple comparisons test showed females spent significantly more time reward seeking during the reward cue than males for sessions R2 (p=0.0274), R3 (p=0.0151) and R5 (p=0.0041). The latency, in seconds, to enter the port post-cue onset was also calculated for each reward cue presentation across all sessions (Figure 1C). Two-way repeated-measures ANOVAs showed a main effect of sex (F(1,34)=20.37, p<.0001), but no significant interaction (F(4,136)=1.684, p=0.1571) or main effect of session (F(4,136)=0.7755, p=0.5429). *Post hoc* Sidak’s multiple comparisons test showed females were significantly faster to enter the port than males during the last 3 reward sessions (R3, p=0.001; R4, p=0.0391; R5, p=0.0014). Taken together, female rats consistently spent more time than males in the reward port during the reward cue in reward pre-training sessions.

**Figure 1.**
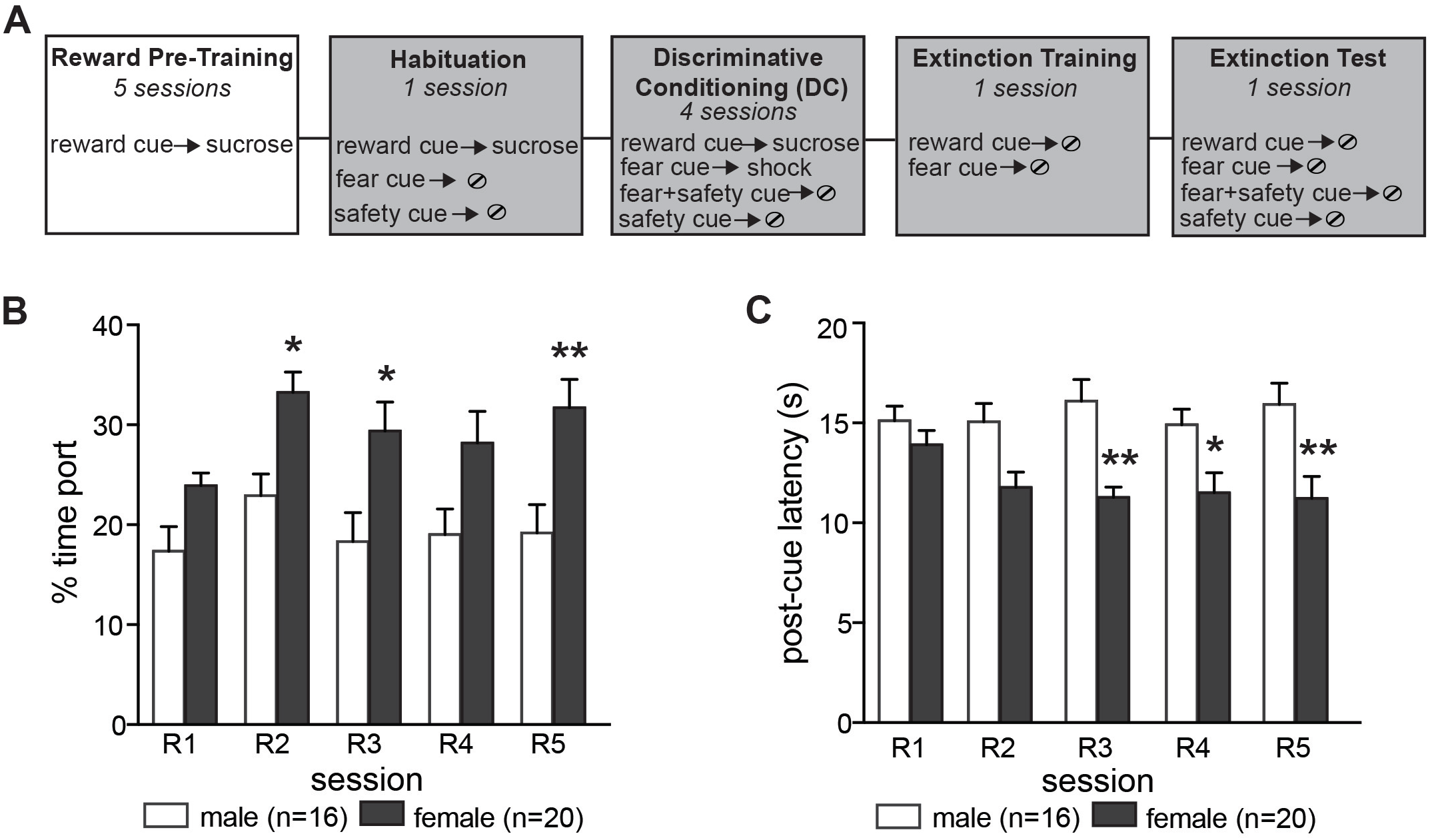
Females show increased reward seeking in response to the reward cue. **A)** Schematic depicting experimental outline. During reward pre-training, rats (16 males, 20 females) received 25 cue-sucrose pairings across 5 separate sessions. **B)** Averaged percent time spent in the reward port during the five reward pre-training sessions (R1-5). Females spent significantly more time in the port compared to males during R2, R3 and R5. **C)** Averaged latency to enter the port after cue onset (in seconds). Females entered the port significantly sooner than males during R3-5. Means +/- SEM. *p<0.05, **p<0.01.

### 3.2 Female rats did not show conditioned inhibition of freezing

After reward pre-training, rats were then exposed to sessions also consisting of reward, fear and safety cues. The reward cue and sucrose reward were the same as the reward pre-training sessions. The fear cue was paired with a 0.5mA footshock, and neither the safety cue nor the fear+safety cue resulted in footshock or sucrose.

The percent time spent at or in the reward port during each cue across session was quantified for each DC session (Figure 2B). Two-way repeated-measures ANOVAs showed a significant cue by sex effect, as well as main effects of cue and sex for DC1 (Table 1). *Post hoc* Tukey’s multiple comparisons test showed that, during DC1, females spent significantly more time reward seeking during the reward cue compared to males (p<0.001), consistent to what was seen in reward pre-training. For the remaining DC2-4 sessions, a main effect of cue was observed (Table 1) and *post hoc* Tukey’s multiple comparisons test showed that both male and female rats spent significantly more time reward seeking during the reward cue compared to all other cues (p<0.0001), with no significant differences between the males and females. Thus, the noticeable increase in reward seeking in the females, that was seen during reward pre-training, dissipated by the 2^nd^ DC session.

**Figure 2.**
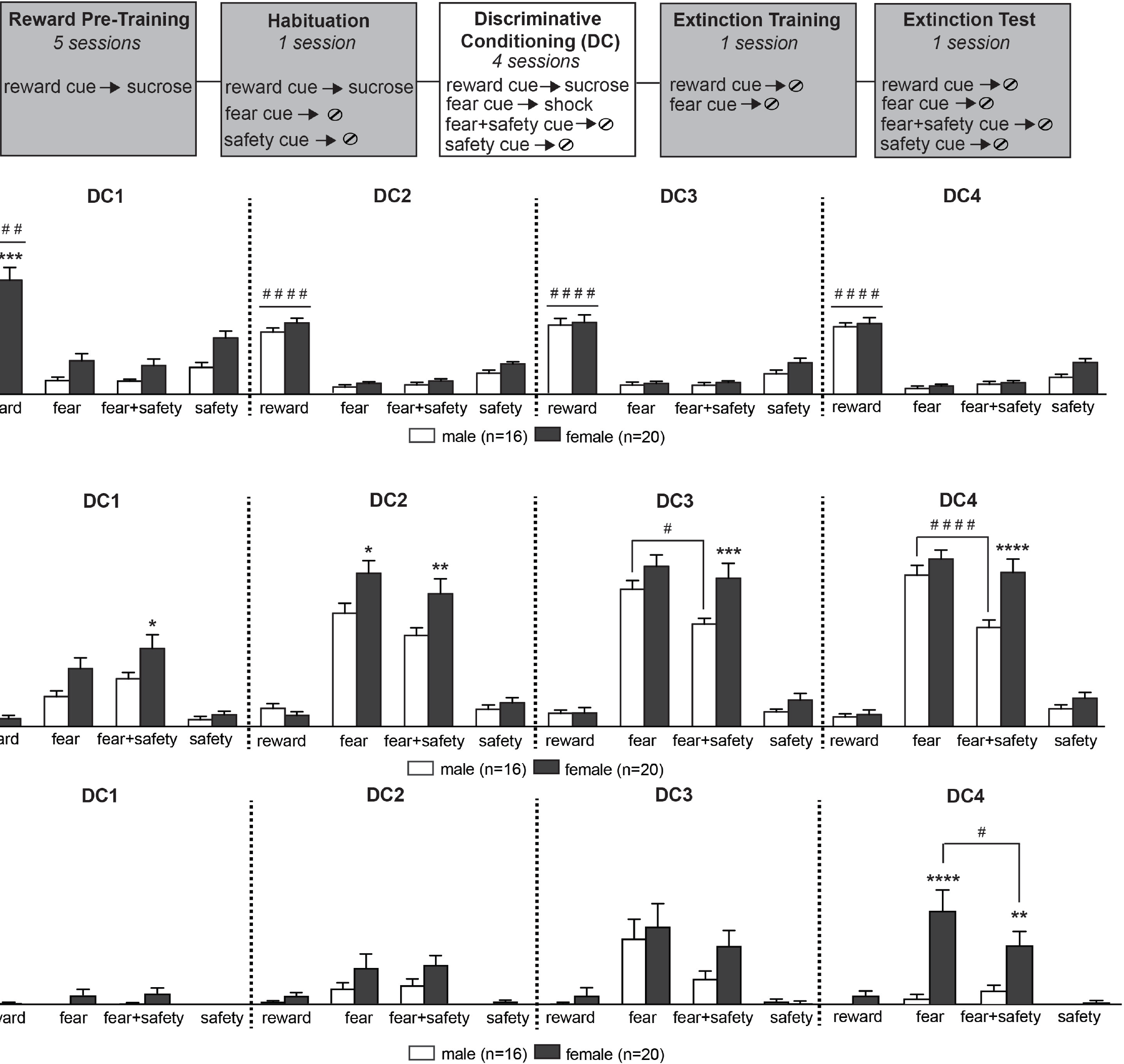
Females do not show inhibition of conditioned freezing in the presence of the safety cue. **A)** Schematic depicting experimental outline. During the 4 DC sessions, rats (16 males, 20 females) were presented with four types of cued trials: reward cue-sucrose, fear cue-shock, fear+safety cue with no footshock and the safety cue presented alone without footshock. **B)** Averaged percent time spent in the port during each cue across the 4 DC sessions. Both males and females showed significantly higher reward seeking during the reward cue compared to all other cues during every DC session. During DC1, females showed significantly higher reward seeking to the reward cue compared to males. **C)** Averaged percent time spent freezing during each cue across the 4 DC sessions. During DC3 and DC4, males showed significantly lower freezing to the fear+safety cue (and reward and safety cues) when compared to the fear cue. Females did not show significant inhibition of conditioned freezing to the fear+safety cue compared to the fear cue during any DC session. Females also showed significantly higher freezing to the fear+safety cue compared to males during every session. **D)** Darting behavior during each cue across the 4 DC sessions. During DC4 females showed significantly more darts than males during the fear and fear+safety cues. Females also showed more darts during the fear cue than the fear+safety cue. Means +/- SEM. # p<0.05, ####p<0.0001 within sex, between cue comparison; * p<0.05, **p<0.01, ***p<0.001, ****p<0.0001 within cue, between sex comparison.

**Table 1.** Summary of two-way repeated-measures ANOVA analyses for reward seeking, freezing and darting behaviors during the four discriminative conditioning (DC) sessions.

| Reward seeking |  |  |  |
| --- | --- | --- | --- |
| *Session* | *Cue x Sex effects* | *Main effect of cue* | *Main effect of sex* |
| DC1 | F(3,102) = 3.472,<br>**p=0.0189** | F(3,102) = 95.16,<br>**p<0.0001** | F(1,34) = 9.827,<br>**p=0.0035** |
| DC2 | F(3,102) = 0.7742,<br>p=0.5110 | F(3,102) = 227.9,<br>**p<0.0001** | F(1,34) = 4.69,<br>**p=0.0374** |
| DC3* | F(3,90) = 0.6512,<br>p=0.5843 | F(3,90) = 117,<br>**p<0.0001** | F(1,30)=1.041,<br>p=0.3157 |
| DC4 | F(3,102) = 2.255,<br>p=0.0864 | F(3,102) = 181.2,<br>**p<0.0001** | F(1,34) = 2.453,<br>p=0.1266 |
| Freezing |  |  |  |
| *Session* | *Cue x Sex effects* | *Main effect of cue* | *Main effect of sex* |
| DC1 | F(3,102) = 2.245,<br>p=0.0876 | F(3,102) = 31.82,<br>**p<0.0001** | F(1,34) = 5.045,<br>**p=0.0313** |
| DC2 | F(3,102) = 4.075,<br>**p=0.0089** | F(3,102) = 103.4,<br>**p<0.0001** | F(1,34) = 6.621,<br>**p=0.0146** |
| DC3* | F(3,90) = 2.9,<br>**p=0.0393** | F(3,90) = 151.3,<br>**p<0.0001** | F(1,30)=9.719,<br>**p=0.0040** |
| DC4 | F(3,102) = 4.889,<br>**p=0.0032** | F(3,102) = 198.9,<br>**p<0.0001** | F(1,34) = 8.294,<br>**p=0.0068** |
| Darting |  |  |  |
| *Session* | *Cue x Sex effects* | *Main effect of cue* | *Main effect of sex* |
| DC1 | F(3,102) = 1.98,<br>p=0.1216 | F(3, 102) = 2.388,<br>p=0.0733 | F(1,34) = 4.146,<br>**p=0.0496** |
| DC2 | F(3,102) = 1.134,<br>p=0.3390 | F(3, 102) = 9.377,<br>**p<0.0001** | F(1,34) = 3.667,<br>p=0.0640 |
| DC3 | F(3,102) = 0.9158,<br>p=0.4361 | F(3, 102) = 18.96,<br>**p<0.0001** | F(1,34)=0.9579,<br>p=0.3346 |
| DC4 | F(3,102) = 10.65,<br>**p<0.0001** | F(3, 102) = 15.65,<br>**p<0.0001** | F(1,34) = 13.34,<br>**p=0.0009** |

\*video files for 4 females were corrupted for this session (n=16 females, 16 males)

The percent time freezing during each cue across session was quantified for each DC session (Figure 2C). Two-way repeated-measures ANOVAs showed a significant cue by sex effect for sessions DC2-4, as well as main effects of cue and sex for every session (Table 1). *Post hoc* Sidak’s multiple comparisons tests showed that, for every session, females displayed significantly more freezing to the fear+safety cue compared to males (DC1, p=0.0313; DC2, p=0.007; DC3, p=0.0007; DC4, p<0.0001). Females also showed significantly higher freezing levels to the fear cue compared to males during DC2 (p=0.0111). Males showed a significant reduction in freezing levels to the fear+safety cue compared to the fear cue during sessions DC3 (p=0.0156) and DC4 (p<0.0001), thus showing significant conditioned inhibition of freezing. Females did not show a significant inhibition of freezing during any session.

The number of darts during each cue was also quantified for each DC session and expressed as the sum of darts across trials for a given cue divided by the number of trials for that cue (Figure 2D; sum of darts/ # trials). Darting behavior during cue presentation was largely absent until DC3 and DC4. Two-way repeated-measures ANOVAs showed a significant cue by sex effect for DC4, as well as main effects of cue, for DC2-4, and sex, for DC1 and DC4 (Table 1). *Post hoc* Sidak’s multiple comparisons test showed that, during DC4, females expressed more darting behavior compared to males during both the fear cue (p<0.0017) and the fear+safety cue (p=0.0079). Additionally, the females significantly reduced their darting behavior during the fear+safety cue compared to the fear cue (post hoc Tukey’s multiple comparisons test, p=0.0166), suggesting some level of conditioned inhibition of darting behavior.

### 3.3 Female rats did not show significant extinction of freezing

The day after the last DC session all rats underwent fear and reward extinction within the same session. That is, both the fear and reward cues were presented within the same training session, without footshocks or sucrose presentations.

During extinction of reward, there was no main effect of reward trial (F(19,646)=1.526, p=0.0704) or sex (F(1,34)=1.31, p=0.2603) and no interaction (F(19,646)=0.8927, p=0.5924); there was also no significant difference between male and female groups for any trial (Figure 3Bi). One day later when rats were re-tested for extinction memory (Figure 3Bii), there was a main effect of cue (2-way RM ANOVA; F(3,102)=134.7, p<0.0001) and sex (F(1,34)=6.217, p=0.0177). *Post hoc* Sidak’s multiple comparisons test showed females had significantly more port activity than males just during the safety cue (p=0.0452), although this difference did not reflect a large increase in port activity as females spent 6.38% +/- 0.86 of the safety cue in the port compared to 2.66% +/- 0.86 in males. Overall, there appeared to be no differences in the ability of males and females to extinguish their reward seeking responses.

**Figure 3.**
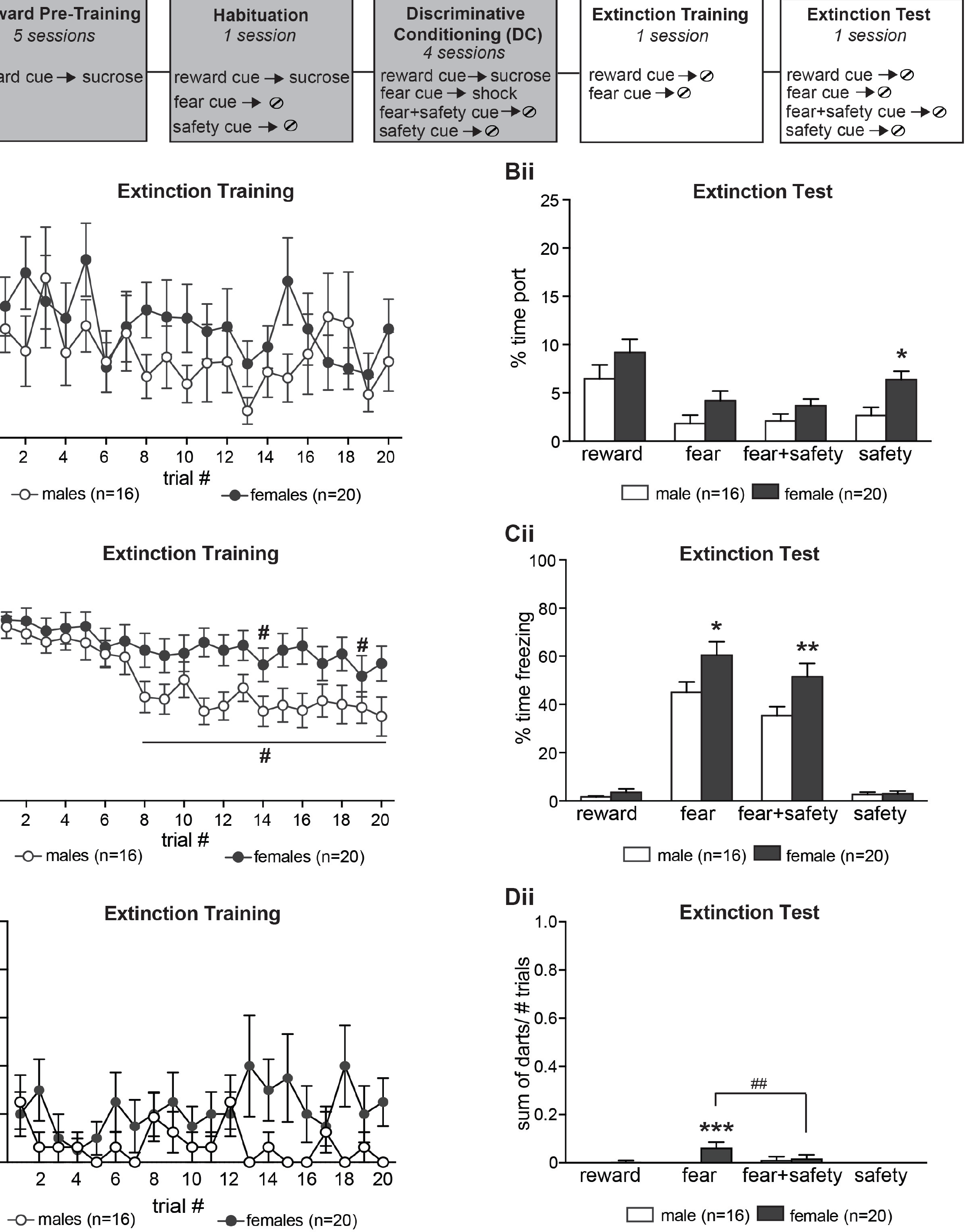
Females do not show significant extinction of fear. A) Schematic depicting experimental outline. During extinction training both the reward and fear cues are presented in the same session without sucrose or footshock. During the test for extinction memory 1 day later all cues are presented without sucrose or footshock. **Bi)** Averaged percent time spent in the port during each reward cue presentation during extinction training. No significant differences were found between males and females during extinction training. **Bii)** Averaged percent time spent in the port during each cue 1 day after extinction training. Females spent significantly more time in the port than males during the safety cue. **Ci)** Averaged percent time spent freezing during each fear cue presentation during extinction training. Compared to the first trial of extinction, males showed significantly reduced freezing during trials 8-20. Freezing levels for females did not significantly decrease at any point in extinction training, with the exception of trials 14 and 19. #p<0.05, compared to trial 1. **Cii)** Averaged percent time spent freezing during each cue 1 day after extinction training. Males showed evidence of fear cue extinction retention. Females froze significantly more than males during the fear and fear+safety cues. **Di)** Averaged darting during each fear cue presentation during extinction training. No significant post hoc differences found between males and females during extinction training. **Dii)** Averaged darting during each cue 1 day after extinction training. Females had significantly higher dart levels than males during the fear cue, which was also significantly higher than the reward, safety and fear+safety cues in females. Means +/- SEM. #p<0.05, ####p<0.0001 within sex, between cue/trial comparisons. *p<0.05, **p<001, ****p<0.0001 within cue, between sex comparisons.

To assess fear extinction the averaged percent time freezing during each trial of fear extinction training was calculated (Figure 3Ci). There was a main effect of fear trial (2-way RM ANOVA; F(19, 646)=7.69, p<0.0001) and sex (2-way RM ANOVA; F(1, 34)=4.607, p=0.0391), but no significant interaction (F(19, 646)=1.566, p=0.059). Compared to trial 1, males showed significantly reduced freezing in extinction trials 8-20 (*post hoc* Dunnett’s multiple comparisons test, p<0.05), demonstrating good fear extinction beginning at the 8^th^ trial. In contrast, females only showed a significant reduction in freezing during trials 14 and 19 compared to the first trial (*post hoc* Dunnett’s multiple comparisons test, p<0.05), demonstrating relatively absent fear extinction. One day later when rats were retested for extinction memory (Figure 3Cii), there was a main effect of cue (F(3,102)=134.7, p<0.0001) and sex (F(1, 34)=6.217, p=0.0177), as well as a significant interaction of cue by sex (F(3, 102)=3.481, p=0.0187). *Post hoc* Sidak’s multiple comparisons test showed that females froze significantly more than males to the fear (p=0.0146) and fear+safety (p=0.0091) cues. This indicates the continued absence of any extinction of freezing in females.

In response to each fear cue presentation across extinction, we also assessed darting levels (Figure 3Di). There was a main effect of sex (F(1,34)=4.816, p=0.0351), but no effect of trial (F(19, 646)=0.6941, p=0.8268) and no significant interaction (F(19, 646)=1.083, p=0.3640). *Post hoc* Sidak’s multiple comparisons test showed no significant differences between males and females for any trial. For the extinction memory test one day later (Figure 3Dii), there was a significant cue by sex interaction (F(3,102)=4.447, p=0.0056), as well as a main effect of both cue (F(3, 102)=4.248, p=0.0072) and sex (F1, 34)=4.834, p=0.0348). Females showed a significantly higher darting levels than males during the fear cue (*post hoc* Sidak’s multiple comparisons test, p=0.0002), which was also significantly higher than the darting levels during the reward (p=0.0002), safety (p<0.0001), and fear+safety (p=0.0082) cues in the females (*post hoc* Tukey’s multiple comparisons test). However, though statistically significant, the amount of darting during the fear cue in females was very low, ranging from 0.05-0.4 across extinction training, and therefore no definitive conclusions can be made regarding darting and extinction in this study.

### 3.4 Shock reactivity in males versus females

To exclude possible sex differences in pain sensitivity and footshock perception, a separate cohort of 8 male and 8 age-matched female rats received 11 unsignaled footshocks of increasing intensities (0.3 mA, 0.35 mA, 0.4 mA, 0.45 mA, 0.5 mA, 0.55 mA, 0.6 mA, 0.7 mA, 0.8 mA, 0.9, 1.0 mA) with an inter-stimulus interval of 2 min. Freezing increased as a function of shock intensities (Figure 4A; 2-way RM ANOVA; F(11,121)=25.9, p<0.0001). No main effects of sex (F(1,121)=0.2871, p=0.6027) or sex by shock (F(11,121)=1.413, p=0.1754) were observed. Our experiments utilized a shock intensity of 0.5mA throughout this study. For this particular intensity, we also noted the number of rats that jumped or darted in response to a 0.5mA shock (Figure 4B,C). No sex differences in the number of rats jumping in response to the 0.5mA footshock were observed (χ^2^: p>0.9). The number of female rats darting after the 0.5mA footshock was higher than males, but not significantly (χ^2^: p =0.0769), with five of the eight female rats tested exhibiting the behavior. A higher number of females darting in response to the footshock in this test would still not explain the lack of conditioned inhibition of freezing in the females, as freezing levels at 0.5mA was slightly lower than the males (Figure 4A). Our results do not definitively show, but do suggest, that females may be more likely to respond to a footshock with a darting response.

**Figure 4.**
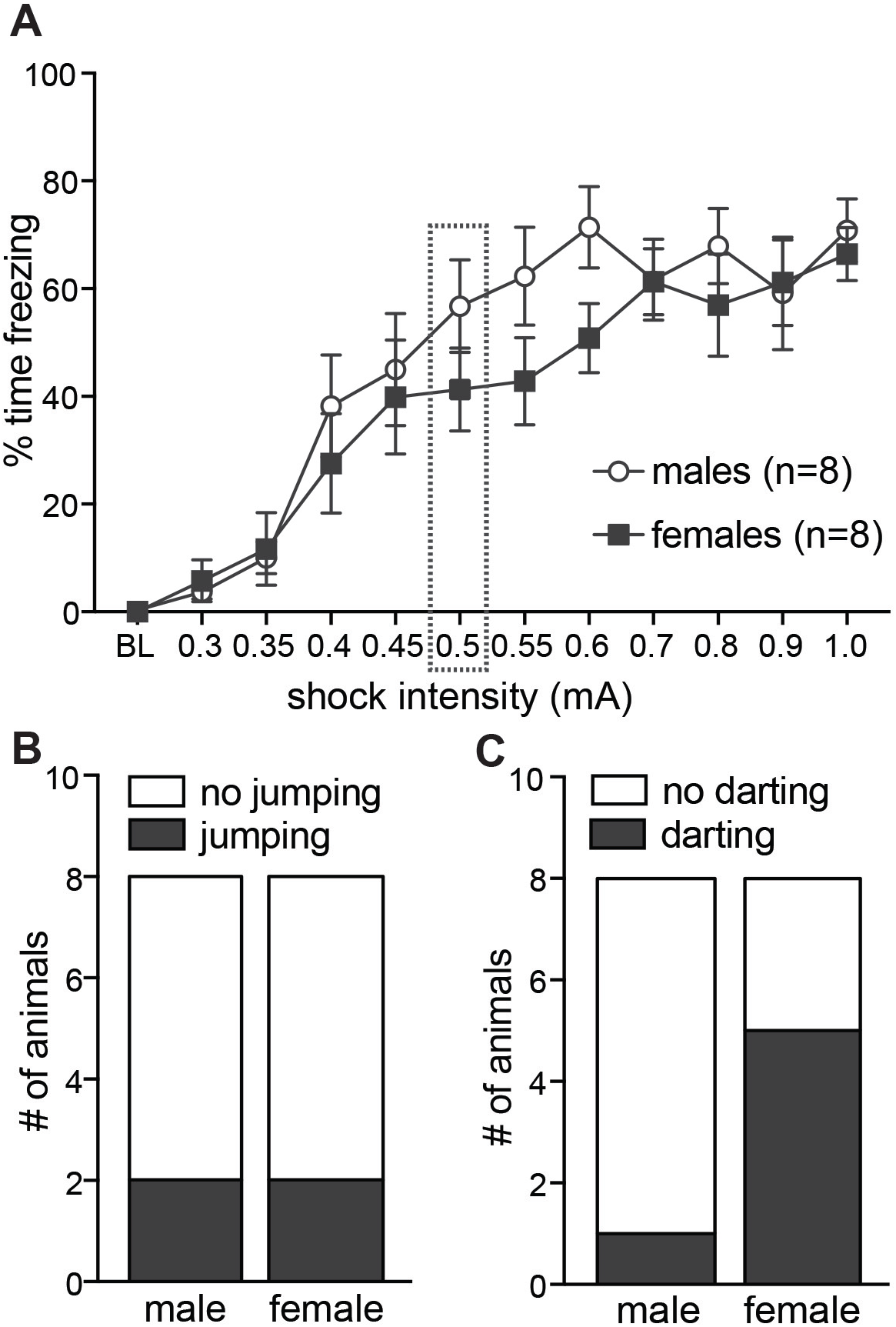
No significant differences in shock reactivity between age-matched male and female rats. A) Male and female rats (n=8 each) were subjected to increasing footshock intensities from 0.3mA to 1.0mA. No significant differences in freezing levels (means +/- SEM) were detected between males and females after each shock presentation. The box around the data at 0.5mA indicates the intensity used for the experiments in this study. There were no significant differences in the number of males or females who jumped **(B)** or darted **(C)** in response to the 0.5mA shock.

## 4. Discussion

In this study, we show females exhibit a significantly different behavioral profile than males in a task that tests for reward, fear and safety cue discrimination, as well as conditioned inhibition and extinction. Female Long Evans rats showed more reward seeking early in training and persistently high freezing levels to the fear cue when in the presence of a safety cue or after fear extinction. Darting behavior in the females late in training showed conditioned inhibition of this behavior in the presence of a safety cue, suggesting the females are able to discriminate between the fear and safety cues but do not suppress their freezing response. This data adds to the growing body of evidence of sex differences in fear regulation and highlights the advantages of using more complex learning paradigms with additional behavioral measurements.

Even though studies including female subjects have been proportionally low, several studies have reported clear sex differences in fear regulation. Most of these are consistent with our findings of reduced discrimination between fear and safety signals. For instance, female mice show more generalization of fear to novel and safe contexts compared to males, and with this generalization there is a concurrent increase in basal amygdala activity (Keiser et al., 2017). Male and female rats also respond differently to the controllability of a stressor. Males display reduced fear during escapable stress versus inescapable stress whereas females exhibit no beneficial effects of perceiving a stressor as escapable and controllable (Baratta et al., 2018). The buffering effects seen in these males were linked to prelimbic cortical neurons projecting to the dorsal raphe nucleus, which do not appear to be engaged in females. Females displaying a similar fear response to both inescapable and escapable stress is similar to our findings of females showing equivalent freezing levels to the fear cue in the presence or absence of a safety cue, in that there were no buffering effects seen by the safety cue. It appears that females do not downregulate their fear response in situations cued as safe.

Our data showing an increase in darting behavior in female rats as the number of fear cue-footshock trials increase is consistent with another report using female rats in a fear conditioning and extinction paradigm (Gruene et al., 2015). Like us, Gruene et al (2015) also show darting levels increase as learning about the fear cue advances. Compared to us, Gruene et al (2015) report notably higher darting frequencies, which is most likely due to the differences in shock intensities and number of trials; our study used 4 trials of 0.5mA per day for 4 days compared to their study using 7 trials of 0.7mA on one day. Our study also includes reinforced reward trials within the same sessions as the fear cue-footshock trials, which could alter the contextual expectations of the training session and reduce overall darting levels. It would be interesting in future studies to identify what leads a female to become a ‘darter’ versus ‘non-darter’. As darting is a more active response compared to freezing, the circuits engaged during potential threats would likely be different in these two populations.

Our findings showing a lack of conditioned inhibition of freezing in females appear to be inconsistent with a recent study demonstrating a lack of sex differences in conditioned inhibition of freezing (Foilb et al., 2018). This is likely due to differences in our respective protocols. First, their footshock intensity was 1.2mA, resulting in freezing levels >90% during the fear cue. As footshock intensity and number of trials are consistently inconsistent across studies, it would be interesting to assess if freezing and darting levels in females follow a linear trend with increasing training intensity, or if there is instead a possibly U-shaped relationship. Foilb et al (2018) also used separate presentations of the fear cue and safety cue throughout training and employed the fear+safety cue summation test during recall, whereas we include fear+safety trials as part of the training. In contrast, another study has shown females discriminate equally to males early in training but then generalize their fear response to the safety cue with continued training (Day et al., 2016). While the females in our study clearly showed equivalent freezing levels to both the fear and fear+safety cues at all time points throughout training, they did not increase their freezing levels to the safety cue when presented alone. And, lastly, our paradigm, unlike others, includes reinforced reward trials during the training of fear and safety cues, which would change the context from a ‘threat-no threat’ situation to a ‘threat-no threat-reward’ situation, inducing approach behaviors on top of defensive behaviors.

Altogether, the data paints a consistent picture of females showing heightened fear responses to cues signaling safety, mimicking the clinical picture in women (Gamwell et al., 2015; Lonsdorf et al., 2015). The presentation of a safety signal not only decreases fear, but also stimulates opposing neuronal activity. Field potential recordings in the striatum during safety signal presentation has shown that brain regions dealing with approach and reward become activated (Rogan et al., 2005). These findings have also been translated to using safety signals to overcome anhedonia in rats (Pollak et al., 2008), showing that safety signals may also be regulating emotion in addition to conditioned behavior (Foilb & Christianson, 2018).

In our study, females consistently showed elevated reward-seeking behavior during the reward cue compared to males beginning in the second reward pre-training session. This data appears consistent with reward studies showing significant sex differences in response to sucrose, with females willing to work more for sucrose in a progressive ratio paradigm (Tapia et al., 2019), and in response to drugs of abuse, with female rats consistently self-administering drugs more rapidly than males (Becker & Koob, 2016). The increased reward-seeking in females seen in our study remained until the end of the first DC session at which point they were equivalent to the males. Interestingly, DC1 is the first time the animals are exposed to footshock. Taking into account the lack of conditioned inhibition of freezing in the females, the females may no longer be as motivated to seek rewards in the face of adverse footshocks. This would be consistent with the report that female rats sacrifice their metabolic needs in order to avoid shocks more than males (Pellman et al., 2017).

Numerous sex differences have been reported in the functioning of the stress neuropeptide, corticotropin-releasing factor (CRF), with differences in receptor expression, distribution, trafficking and signaling (reviewed in (Bangasser & Wiersielis, 2018)). The majority of these differences lead to enhanced CRF efficacy in females, which may lead to heightened sensitivity to stressors in females. Recently, the gene for CRH receptor 1 (*CRHR1*) has been identified as a possible candidate gene for mood and anxiety disorders. Weber et al. (2016) have shown that carrying the *CRHR1* minor rs17689918 allele increases the risk for panic disorders in women. Patients carrying this risk allele also demonstrate more generalization of fear to a safety cue, increased amygdala activation during the safety cue and decreased frontal cortex activation with discriminative fear conditioning. Thus, aberrant CRF signaling can lead to sustained fear under conditions cued as safe and can be manifested by changes in neural activity in the amygdala and frontal cortex.

Neural activity in the amygdala and prefrontal cortex has been shown by our lab to also play a critical role in effective discriminative conditioning in male rats. We have previously identified neurons in the basolateral amygdala (BLA) that discriminate among safety, fear and reward cues in male rats (Sangha et al., 2013); our future experiments will test if females show the same discriminative neurons. Using reversible pharmacological inactivations in male rats, we have also demonstrated that the infralimbic prefrontal cortex (IL) is necessary for suppression of conditioned fear during a safety cue and the prelimbic prefrontal cortex (PL) is necessary for fear expression and discriminatory reward-seeking (Sangha, Robinson, et al., 2014). These results indicate that activating the IL in the females may improve conditioned inhibition to the combined fear and safety cues. Our results with male rats also show that manipulating D1-receptor mediated dopamine activity in the BLA disrupts suppression of conditioned fear (Ng et al., 2018), implicating dopaminergic ventral tegmental area (VTA) neurons projecting to the BLA in safety-fear-reward discrimination.

Our findings are consistent with human studies where females show less discrimination between the fear and safety signals than males (Gamwell et al., 2015; Lonsdorf et al., 2015), which may reflect underlying mechanisms of increased prevalence for anxiety and stress-related disorders in women. For example, a deficiency in effective safety signal processing has been linked to Post-traumatic Stress Disorder (Jovanovic et al., 2009, 2010), panic disorder (Gorka et al., 2014), and anxiety (Lissek et al., 2005), all disorders with a higher incidence in women than men (Mclean et al., 2011). In our paradigm, females show a significantly different behavioral profile than males that is consistent with the clinical picture, thus making it a great tool to test for the neurobiological mechanisms underlying these sex differences.

## 5. Acknowledgements

We thank Yolanda Jonker, Signe Hobaugh, Jessica Kerns, Emma Speckelson, Emily Willis, and Mackenzie McIntosh for excellent animal care. IM was supported by the Alexander von Humboldt foundation. This work was supported by the National Institutes of Health [NIMH R01MH110425 to SS].

